# Lnc-*ECAL-1* controls cerebrovascular homeostasis by targeting endothelium-specific tight junction protein *Cldn5b*

**DOI:** 10.1101/2020.01.30.926279

**Authors:** Fang-Fang Li, Yu-Lai Liang, Jing-Jing Zhang, Qing Jing

## Abstract

Cerebrovascular disorder-induced brain blood flow interruption or intracranial hemorrhage pose a great threaten to health. Emerging roles of long-noncoding RNAs (lncRNAs) in diagnosis and treatment of cardiovascular diseases have been recognized. However, whether and how lncRNAs modulate vascular homeostasis, especially network formation remain largely unknown. Here, we identified *ECAL-1*, a long non-coding RNA, as an important determinant for cerebrovascular homeostasis. Using the morpholino- and CRISPR /Cas9-based genetic modifications in combination with *in vivo* confocal imaging in zebrafish, we claimed that inactivation of *ECAL-1* induced the apparent distortion of cerebral vascular pattern accompanied by intracranial hemorrhage. These cerebrovascular abnormalities were associated with decreased proliferation and anomalous interconnection of endothelial cells. Importantly, overexpression of Cldn5b, an endothelial cell-specific tight junction protein-encoding gene, could partially rescued the phenotype induced by *ECAL-1* deficiency. Furthermore, bioinformatic analysis and experimental validation revealed that *ECAL-1* sponged miR-23a, which targeted Cldn5b 3’UTR and modulated Cldn5b expression, to maintain cerebrovascular pattern formation and integrity. Our results presented here revealed that *ECAL-1* specifically controls cerebrovascular network formation and integrity through targeting miR-23a-*Cldn5b* axis. These findings provide a new regulation modality for cerebrovascular patterning and the potential neurovascular disorders, and *ECAL-1*-miR-23a axis represents as an attractive therapeutic target for cerebrovascular diseases.

## Introduction

The homeostasis of brain vascular network is essential for the maintenance of neuronal activities and the relevant physiological functions, and its disorders result in severe pathologies, such as multiple sclerosis, hemorrhagic stroke, epilepsy etc. (Obermeier et al., 2013). Indeed, vascular development undergoes two sequential stages, i.e., vasculogenesis and angiogenesis. Following the formation of vascular network, blood vessels proceed to reshape through selectively fusing and degenerating, and then recruiting peripheral cells, i.e., pericytes, astrocytes, and microglia, to form mature blood vessels (Carmeliet and Jain, 2011; Geudens and Gerhardt, 2011). In the brain, endothelial cells (ECs) are specialized and particularly important in these processes (Dejana et al., 2009; Obermeier et al., 2013), despite both ECs connection and pericytes are involved in the maintenance of vascular integrity (Armulik et al., 2010; Nitta et al., 2003; Wang et al., 2014; Weis and Cheresh, 2005; Xu et al., 2017).

Non-coding RNAs emerge as important regulators and effectors in cardiovascular development and diseases (Fish et al., 2008; Wang et al., 2008; Xu et al., 2017; Zhou et al., 2014; Zou et al., 2011). Long non-coding RNAs (lncRNAs) are non-protein coding transcripts longer than 200 nucleotides, and express in a tissue-specific manner (Orom et al., 2010; Rinn and Chang, 2012). They can act as signaling, decoying, guiding or scaffolding molecules to mediate pathophysiological functions (Rinn and Chang, 2012). For instance, *Tie-1 AS* is essential for EC contact junctions by selectively binding to *Tie-1* mRNA (Li et al., 2010). *Braveheart* functions upstream of mesoderm posterior 1 (MesP1) and contributes to cardiovascular lineage commitment (Klattenhoff et al., 2013). *STEEL* modulates angiogenic behavior by forming a complex with poly(ADP-ribose) polymerase 1 (PARP1) (Man et al., 2018). *LincRNA-p21* represents a key regulator of cell proliferation and apoptosis during atherosclerosis (Wu et al., 2014), and sponges miR-130b to promote EC apoptosis and cell cycle progression (He et al., 2015). However, the roles of lncRNAs in organ-specific endothelial function and vascular homeostasis remain unclear.

Hundreds of lncRNAs have been identified in zebrafish (Kaushik et al., 2013; Pauli et al., 2012; Ulitsky et al., 2011). Given the great convenience of brain vascular network visualization and genetic manipulation (Chen et al., 2012; Fujita et al., 2011; Vanhollebeke et al., 2015; Xu et al., 2017), herein we adopted zebrafish to elucidate the roles of lncRNAs in cerebrovascular homeostasis and the underlying molecular mechanisms. Here, we identified a lncRNA as endothelial connection associated lncRNA (*ECAL-1*) in zebrafish, and claimed that *ECAL-1* was indispensable for brain EC connection, and determined EC proliferation *in vivo* and central arteries (CtAs) morphogenesis.

## Results

### *ECAL-1* is required for cerebrovascular homeostasis in zebrafish

To explore the *in vivo* function of *ECAL-1*, we adopted morpholino- and CRISPR/Cas9-mediated gene modifications in zebrafish. The morpholino knockdown was carried out in wild type zebrafish embryos. *ECAL*-*1* MO was designed to target the splicing site of *ECAL-1*, and micropeptide (MP) MO inhibited the translation of micropeptide (87 amino acids) in *ECAL-1* (Fig.1A). We found that *ECAL-1* MO reduced the expression of *ECAL-1* (Fig. S1D), and MP MO downregulated the expression of GFP bearing a binding site of MP MO (Fig. S1A and S1B). In addition, the expression of *p53* were not changed in embryos injected with *ECAL-1* morpholino (1 pmol) (Fig.S1C), precluding the morpholino-mediated off-target effects that may lead to excessive activation of *p53* (Robu et al., 2007; Rossi et al., 2015). With this approach, we observed that morphology of embryos injected with *ECAL-1* MO and MP MO appeared normal, while ~20% of *ECAL-1* morphants displayed intracranial hemorrhage at 72 hours post fertilization (hpf) (Fig.1C and 1D).

**Figure 1.**
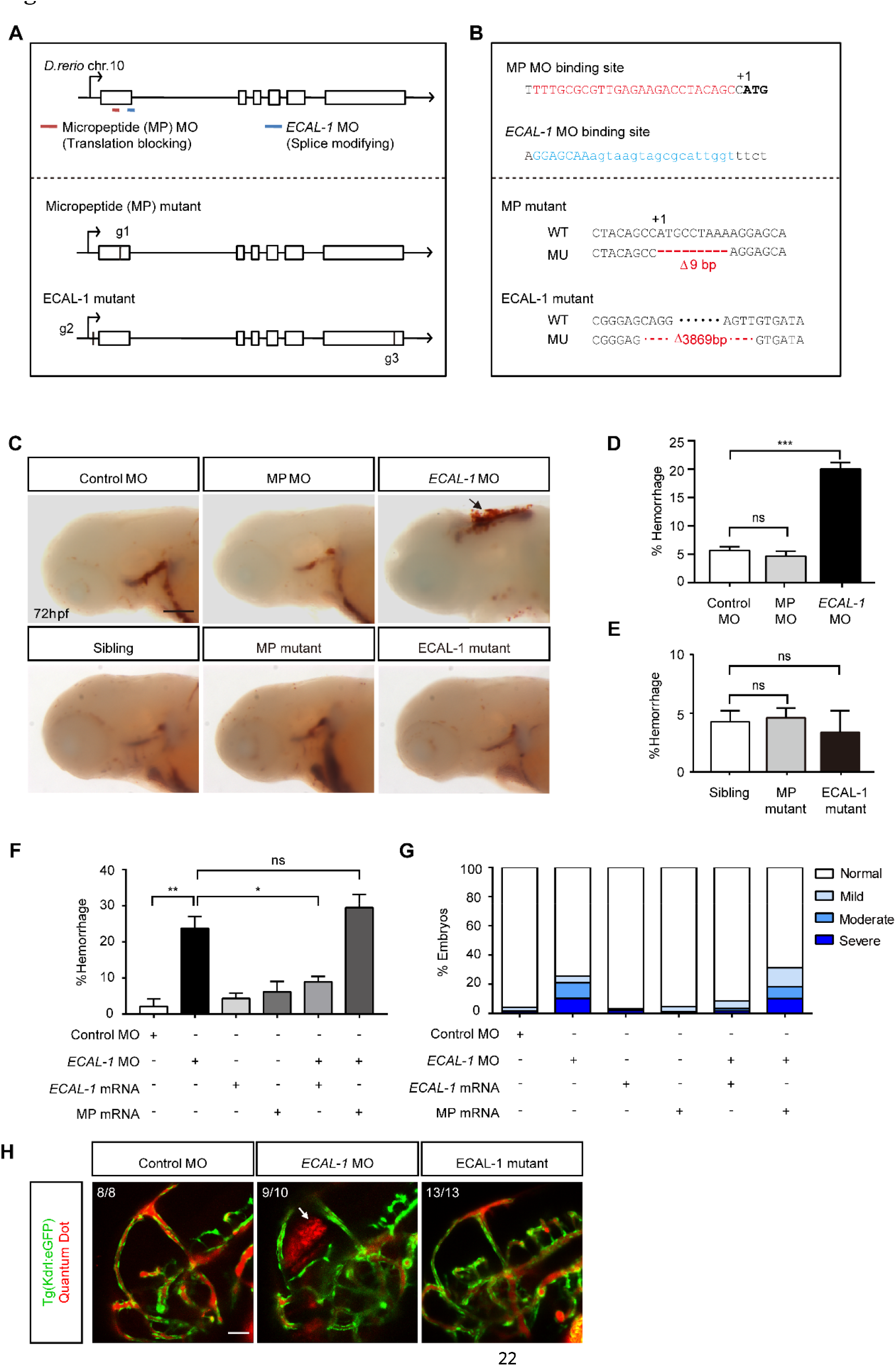
*ECAL-1* was essential for cerebrovascular homeostasis. **(A)** Diagram and sequence of morphorlinos (MO) target sites. **(B)** Schematic diagram of ECAL-1 knockout line and the mutated sequences of micropeptide (MP) mutant and ECAL-1 mutant. **(C)** Bright field images of O-dianisidine staining embryos, including morphants and mutants. Hemorrhage sites are indicated by black arrows. **(D and E)** Statistics of embryos with cranial hemorrhage in *ECAL-1* morphants and mutants. (D) N=304, 303 and 320 for control MO, MP MO and *ECAL-1* MO, respectively. (E) N=387, 501 and 151 for Sibling, MP mutant and ECAL-1 mutant, respectively. Statistical analysis was conducted using one-way ANOVA *post hoc* Tukey test. **(F)** Statistics of cranial hemorrhage in embryos injected with control MO, *ECAL-1* MO, *ECAL-1* mRNA, micropeptide (MP) mRNA, (*ECAL-1* MO and *ECAL-1* mRNA), (*ECAL-1* MO and MP mRNA). Statistical analysis was conducted using one-way ANOVA *post hoc* Tukey test. **(G)** Percentage of cranial hemorrhage in embryos injected with control MO, *ECAL-1* MO, *ECAL-1* mRNA, MP mRNA, (*ECAL-1* MO and *ECAL-1* mRNA), (*ECAL-1* MO and MP mRNA) in different degree. (F and G) N=127, 117, 103, 86, 89 and 99 for control MO, *ECAL-1* MO, *ECAL-1* mRNA, (*ECAL-1* MO + *ECAL-1* mRNA) and (*ECAL-1* MO + MP mRNA), respectively. **(H)** Confocal stack micrographs of head vasculature injected with Quantum Dot at 55hpf, including embryos injected with control embryos, morphants and mutants, respectively. Leakage sites are indicated by white arrows. N+8, 10 and 13 for control MO, *ECAL-1* MO and ECAL-1 mutant, respectively. Values are means□±□SEM. *p<0.05; ***p<0.001. Scale bar: 200μm (C); 100μm (H).

Based on the CRISPR/Cas9 system, we generated two knockout lines, i.e., MP mutant without micropeptide translation, and ECAL-1 mutant without *ECAL-1* expression (Fig.1B and S1E). The expression of *ECAL-1* was reduced to 40% in MP mutants, and we detected barely no expression of *ECAL-1* in ECAL-1 mutant (Fig.S1E). Approximately 5% of embryos exhibited intracranial hemorrhage, which was no significantly difference among siblings, MP mutants and ECAL-1 mutants (Fig.1C and 1E).

Then, to validate the functional phenotype by *ECAL-1* knockdown, we adopted the full length of *ECAL-1* and the coding sequence of micropeptide. Replenishment of *ECAL-1* capped mRNA, rather than the coding sequence of micropeptide, decreased the hemorrhage rate in *ECAL-1* morphants (Fig.1F and 1G), supporting the role of *ECAL-1* in cerebrovascular homeostasis. Experiments with Quantum Dot injection further demonstrated the disruption of cerebral vessels in *ECAL-1* morphants (Fig.1H), highlighting the importance of *ECAL-1* in the development of cerebrovascular network.

Collectively, *ECAL-1* is responsible for cerebrovascular integrity to maintain homeostasis during development.

### *ECAL-1* determines pattern formation of cerebrovascular network

To further define the vascular abnormalities associated with cerebrovascular homeostasis, we traced the dynamic changes of cerebrovascular network using the transgenic zebrafish *Tg(Kdrl:eGFP)*. The central arteries (CtAs) sprout from PHBC from 32 hpf to 48 hpf (Fujita et al., 2011; Vanhollebeke et al., 2015). Notably, we observed that hemorrhage occurred from 36 hpf (Data not shown), and most hemorrhage events occurred from 48 hpf to 60 hpf and appeared more severe from 60 hpf to 72 hpf (Fig.2A). The occurrence of intracranial hemorrhage was coincident with the time window of cerebrovascular network formation, further underscoring the indispensability for *ECAL-1* in cerebrovascular development.

**Figure 2.**
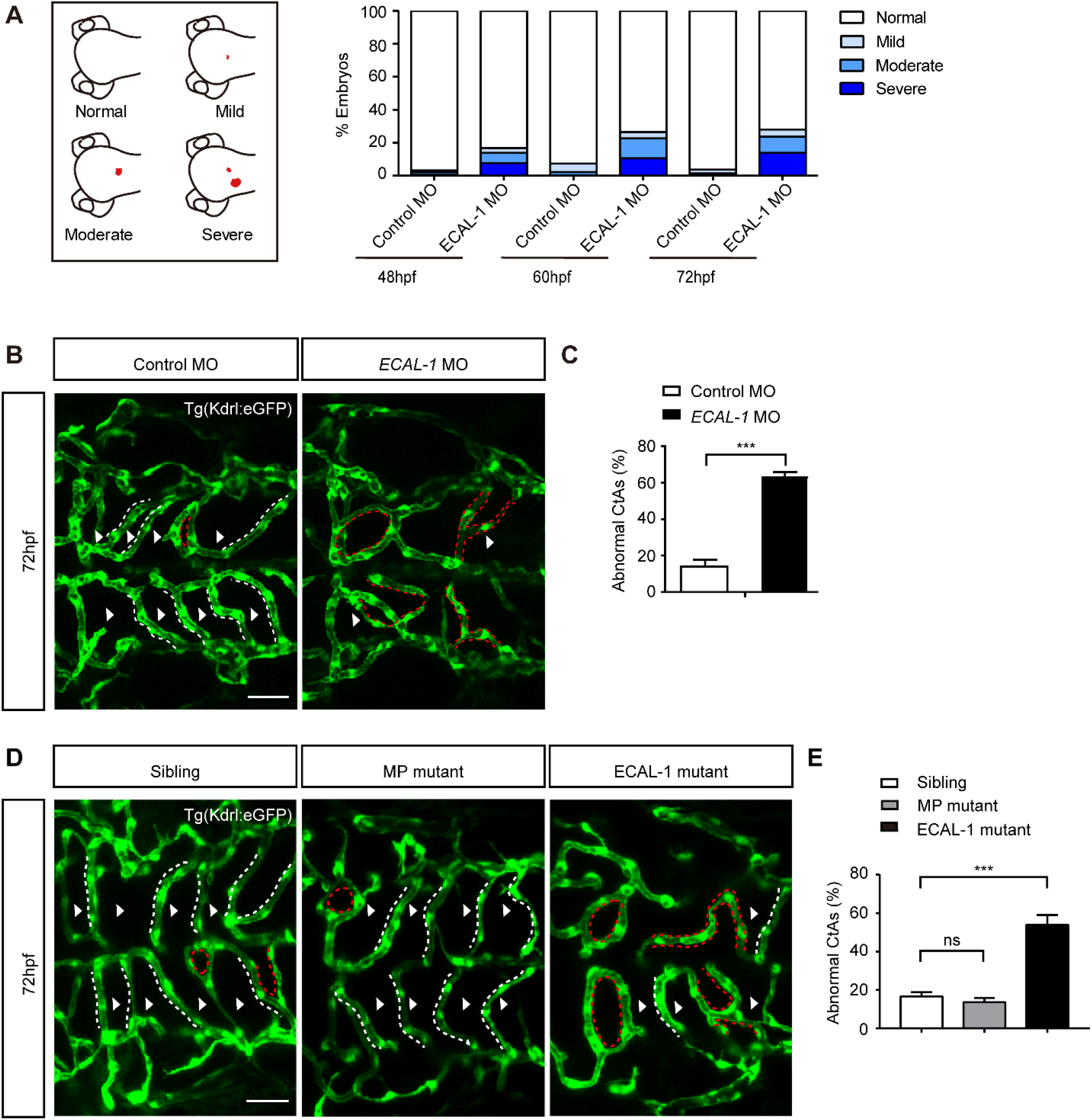
*ECAL-1* determines pattern formation of cerebrovascular network. **(A)** Percentage of cranial hemorrhage in control embryos and *ECAL-1* morphants in different degree at 48hpf, 60hpf and 72hpf. N=353 and 339 for control MO and *ECAL-1* MO, respectively. **(B)** Confocal stack micrographs of embryos injected with control MO and ECAL-1 MO. Dorsal view of hindbrain at 72hpf. **(C)** Quantification percentage of abnormal hindbrain CtAs of control embryos and *ECAL-1* morphants at 72hpf. N=9 and 9 for control MO and *ECAL-1* MO, respectively. Statistical analysis was conducted using unpaired Student’s two-tailed *t*-test. **(D)** Confocal stack micrographs of Sibling, MP mutant and ECAL-1 mutant. Dorsal view of hindbrain at 72hpf. Statistical analysis was conducted using one-way ANOVA *post hoc* Tukey test. **(E)** Percentage of abnormal CtAs in Sibling, MP mutant and ECAL-1 mutants at 72hpf. The abnormal CtAs are indicated by red dotted line, and normal CtAs are highlighted by white dotted line. The white arrows indicate the right cerebrovascular patterning. N=13, 11 and 12 for Sibling, MP mutant and ECAL-1 mutant, respectively. Values are means□±□SEM. ***p<0.001; Scale bar: 50μm (B and D).

In *ECAL-1* morphants and mutants, both the number of CtAs (Fig.S2A, S2B, S2D and S2E) and the penetration depth of CtAs into the hindbrain matter (Fig.S2A, S2C, S2D and S2F) were not affected. However, the pattern of CtAs displayed pruning structures (“H” and “O” type) and anomalous connection in *ECAL-1* morphants and ECAL-1 mutants (Fig.2B-2E), but not in MP mutants (Fig. 2D and 2E). The morphology of primary vessels in brain (PHBC, MCeV, LDA and CCV) (Fig.S3A-S3D) and trunk (ISV and DLAV) (Fig.S3E and S3F) remained intact in *ECAL-1* morphants. Together, *ECAL-1* modulates the pattern formation of CtAs.

### *ECAL-1* is expressed in endothelial cells and neuron of brain

To dissect the action modality of *ECAL-1* in regulating cerebrovascular development, we characterized its expression pattern. Using whole mount *in situ* hybridization, we identified that *ECAL-1* was distributed in blood vessels and parenchyma of brain (Fig.3A c and d). Frozen sections highlighted the expression of *ECAL-1* in vessel wall and the inner part of brain (Fig.3A h and i). *ECAL-1* expression remained at a high level until 120 hpf (Fig.3B). FACS assay confirmed the expression of *ECAL-1* in endothelial cells, while highly in non-endothelial cells (Fig.3C). Fluorescence *in situ* hybridization (FISH) results showed that *ECAL-1* was expressed in some of *flk1*-positive endothelial cells in trunk at 28 hpf, more was expressed in dorsal nervous system (Fig.3D). At 48 hpf, *ECAL-1* was mainly co-localized with neuron marker *HuC* in head (Fig.3E).

**Figure 3.**
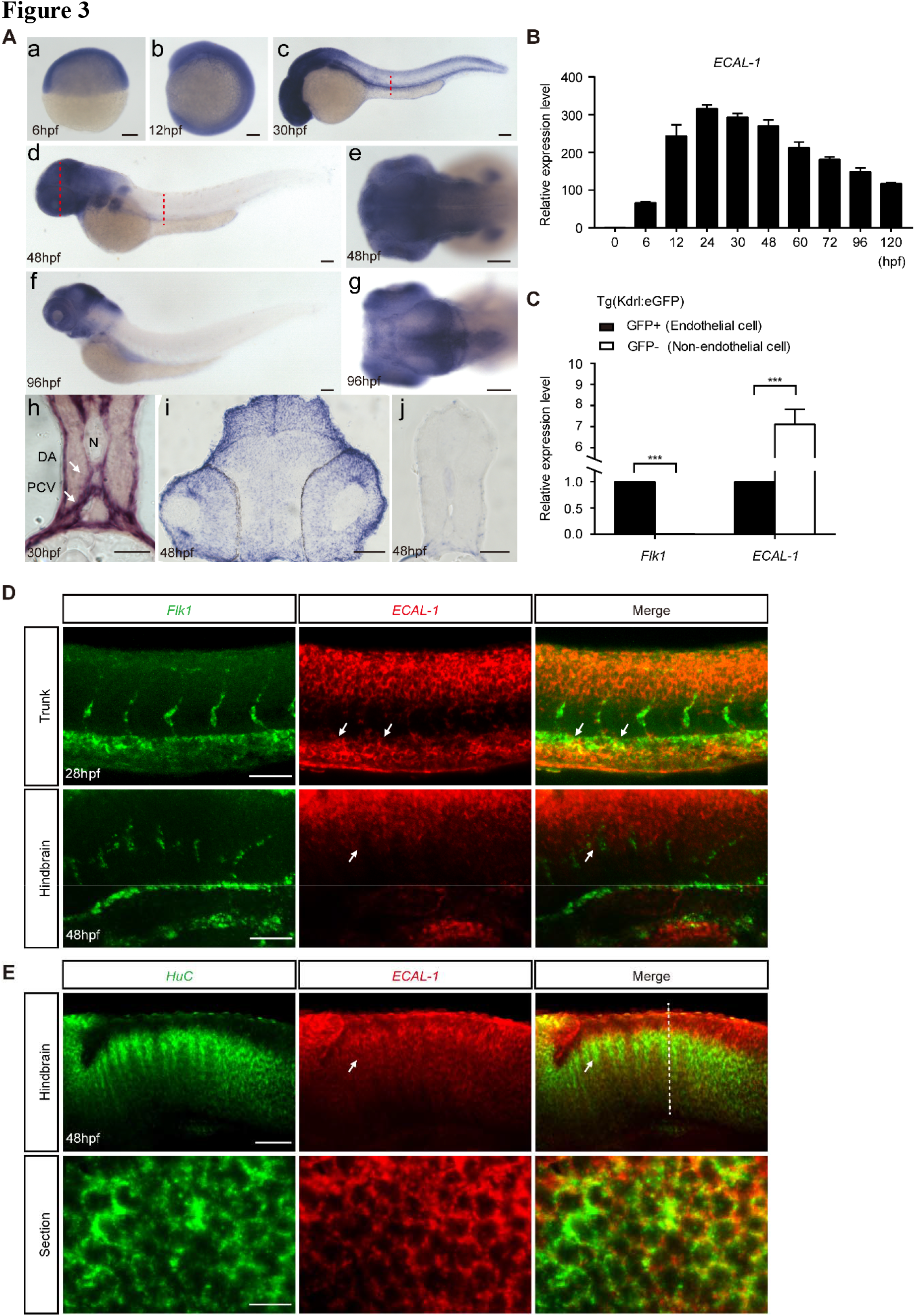
*ECAL-1* is enriched in endothelial cells and neurons of brain. **(A)** Expression pattern of *ECAL-1*. (a-g) Whole mount *in situ* hybridization (WISH) in indicated embryos. (h-j) Frozen section for trunk and head. DA: Dorsal aorta, PCV: Posterior cardinal vein, N: notochord. White arrows indicate co-localization of ECAL-1 and marker genes (*Flk1* or *HuC*). Anterior is to the left except as noted. **(B)** Quantification of *ECAL-1* expression level in different developmental stages by Real-Time PCR. **(C)** Quantification of *ECAL-1* expression level at FACS-sorted vascular endothelial cells (GFP positive) and non-endothelial cells (GFP negative) using Tg(Kdrl:eGFP) transgenic embryos. Each group has about 100 embryos, and this assay was performed three times. Statistical analysis was conducted using unpaired Student’s two-tailed *t*-test. **(D)** Double fluorescence *in situ* hybridization (DFISH) for *Flk1* and *ECAL-1* in trunk and hindbrain. **(E)** Whole-mount and section double fluorescence in situ hybridization for *HuC* and *ECAL-1* in hindbrain. The embryos used in Values are means□±□SEM. ***p<0.001. Scale bar: 100μm (a-g in A); 50μm (h-j in A); 50μm (D and top row in E). 10μm (bottom row in E).

### *ECAL-1* defect disrupts the connection between endothelial cells

We further asked whether *ECAL-1* modulated the biology of endothelial cells. During this process, migration and proliferation are the main behaviors of vascular endothelial cells. Through measuring the depth of CtAs penetrated into brain matter, we found that *ECAL-1* deficiency didn’t affect the migration of ECs (Fig.S2A and S2C). However, the ECs numbers in CtAs was decreased in the *Tg*(*Fli1:negfp,Kdrl:mcherry*) embryos with *ECAL-1* deficiency (Fig.4A and 4B), and those in PHBC was comparable with sibling controls (Fig.4A and 4C). To define the potential role of pericytes in cerebrovascular abnormalities by *ECAL*-1 deficiency, we detected the expression of *Pdgfrb*, a pericyte marker. As shown in Fig.4D and 4E, the pericyte coverage of hindbrain CtAs remained unaffected in *ECAL-1* morphants, precluding the contribution of pericytes. Ultra-structure analysis of ECs showed that intercellular junctions were discontinuous. Particularly, intercellular cleft was widened in *ECAL-1* morphants with intracranial hemorrhage (Fig.4F). These results indicated that *ECAL-1* is essential for EC proliferation and intercellular connection in CtAs.

**Figure 4.**
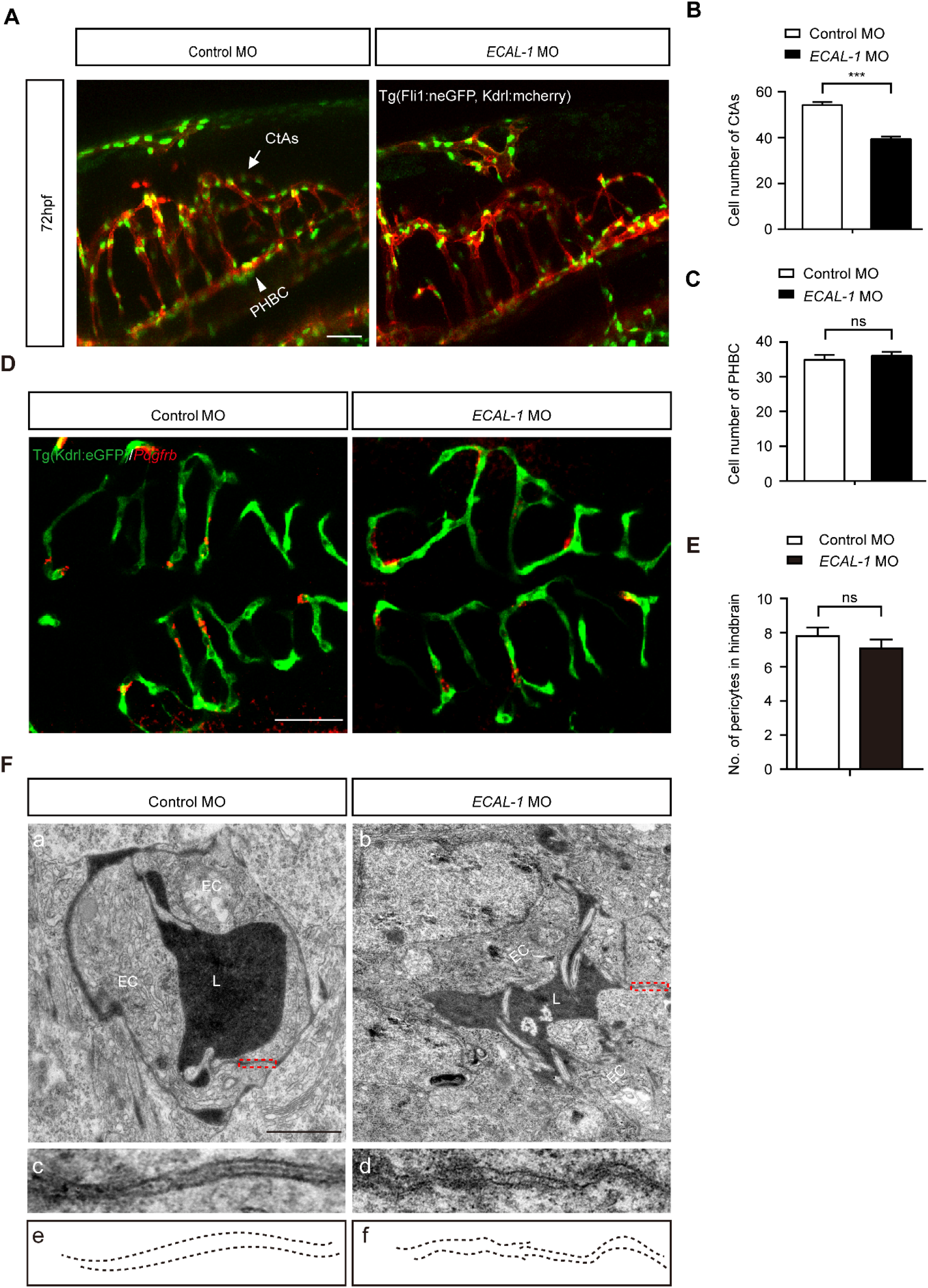
Connections between endothelial cells were disturbed in ECAL-1 deficiency embryos. **(A)** Lateral view of head vasculature in Tg(*Fli1:negfp,Kdrl:mcherry^Ras^*) embryos injected with control MO and *ECAL-1* MO at 72hpf. **(B and C)** Statistics of endothelial cell number in PHBC and CtAs (C) of control embryos and *ECAL-1* morphants at 72hpf. N-8 and 10 for control MO and *ECAL-1* MO, respectively. Statistical analysis was conducted using unpaired Student’s two-tailed *t*-test. **(D)** Fluorescence *in situ* hybridization for *Pdgfrb* and immunofluorescence for GFP in control embryos, *ECAL-1* morphants and ECAL-1 mutants. **(E)** Statistics of pericytes in hindbrain of control embryos, *ECAL-1* morphants. Statistical analysis was conducted using one-way ANOVA *post hoc* Tukey test. N-8 and 8, for control MO, *ECAL-1* MO, respectively. **(F)** Transmission electron microscopy (TEM) images showing the ultrastructural changes of brain vessels in control embryos and *ECAL-1* morphants. (a and b) The cross section of CtAs. (c and d)The areas outlined by red dashed frame in (a and b). (e and f) Schematic diagram of endothelial cell connection. L:Lumen; EC: Endothelial cell; Values are means□±□SEM. ***p<0.001. Scale bar: 50μm (A); 50μm (D); 1μm (a and b in F).

### *ECAL-1* controls the expression of endothelial tight junction protein Cldn5b to affect cerebrovascular homeostasis

The maintenance of EC connection is predominated by intercellular junctional complexes comprised of tight junction molecules, e.g., adhesion protein Cdh5 and Cldn5 (Dejana et al., 2009; Obermeier et al., 2013). We checked the expression of these tight junction-related molecules, with the results that Cldn5b was reduced, while Cdh5 appeared normal in *ECAL-1* morphants at 30 hpf (Fig.5A). Western blotting examination demonstrated that Cldn5 protein was significantly decreased in *ECAL-1* morphants, and Cdh5 protein was comparable with that in sibling controls at 48 hpf (Fig.5B and 5C). Moreover, immunofluorescent imaging revealed the expression of *Cldn5* in brain vessels, and validated the reduction of Cldn5 proteins by *ECAL-1* deficiency (Fig.5D).

**Figure 5.**
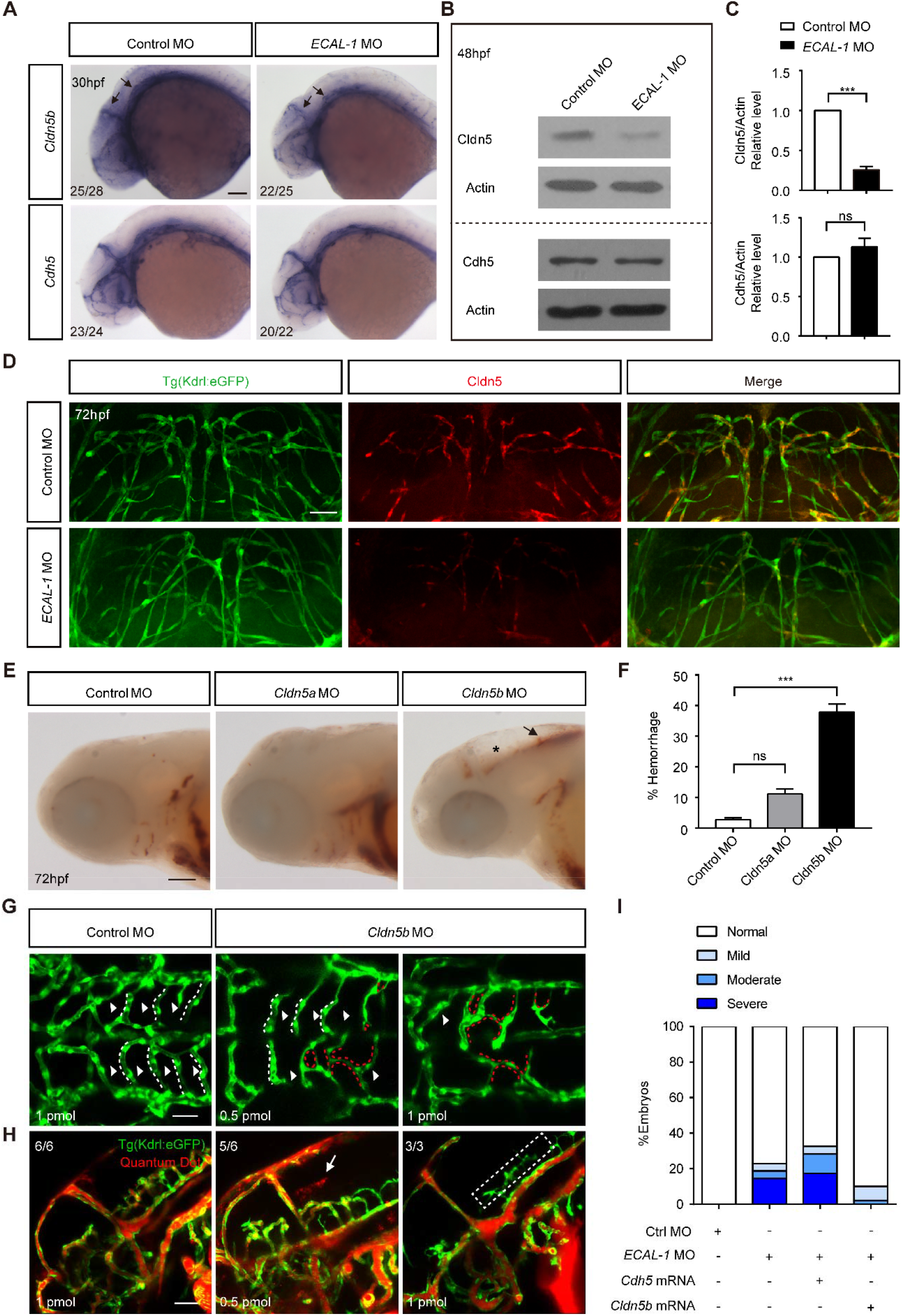
Cldn5b is a functional downstream effector of *ECAL-1* during cerebrovascular homeostasis maintenance. **(A)** Detection of *Cldn5b* and Cdh5 mRNA expression level by whole mount in situ hybridization in control embryos and *ECAL-1* morphants at 30hpf. **(B)** Detection of Cldn5 and Cdh5 protein expression level by Western Blotting at 48hpf. Each group has more than 30 embryos, and this assay was performed three times. **(C)** Quantitative analysis of three times western blotting results. **(D)**Immunofluorescence for GFP and Cldn5 in cross section of control embryos and *ECAL-1* morphants at 72hpf. **(E)** Bright field images of O-dianisidine staining embryos injected with control MO, *Cldn5a* MO or *Cldn5b* MO at 72 hpf. **(F)** Statistics of embryos with cranial hemorrhage, including control embryos, *Cldn5a* morphants and *Cldn5b* morphants at 72hpf. N=168, 108 and 153, for control MO, *Cldn5a* MO and *Cldn5b* MO, respectively. Statistical analysis was conducted using one-way ANOVA *post hoc* Tukey test. **(G)** Confocal stack micrographs of embryos injected with control MO and *Cldn5b* MO (0.5pmol and 1pmol). Dorsal view of hindbrain at 72hpf. The abnormal CtAs are indicated by red dotted line, and normal CtAs are highlighted by white dotted line. The white arrows indicate the right cerebrovascular patterning. **(H)** Confocal stack micrographs of head vasculature injected with Quantum Dot, including control embryos, *Cldn5b* morphants (0.5 pmol and 1 pmol). **(I)** Percentage of cranial hemorrhage in embryos injected with control MO, *ECAL-1* MO, (*ECAL-1* MO and *Cdh5* mRNA), or (*ECAL-1* MO and *Cldn5b* mRNA) in different degree. The cranial edema is indicated by asterisk. N=121, 155, 140 and 152, for control MO, *ECAL-1* MO, (*ECAL-1* MO + *Cdh5* mRNA) and (*ECAL-1* MO + *Cldn5b* mRNA), respectively. Values are means□±□SEM. *p<0.05; ***p<0.001. Scale bar: 100μm (A); 25μm (D); 200μm (E); 50μm (G); 100μm (H).

CLDN5 plays a vital role in blood brain barrier integrity (Nitta et al., 2003; Wang et al., 2012), and its role in cerebrovascular homeostasis deserves further study. In zebrafish, *Cldn5* has two orthologous genes, *Cldn5a* and *Cldn5b* (Xie et al., 2010). To explore the role of Cldn5 in cerebrovascular homeostasis, we designed MOs targeting the ATG start site of *Cldn5a* or *Cldn5b*, respectively (Fig.S4A). As shown in Fig.S4B-S4E, the level of Cldn5 protein was reduced in *Cldn5a* morphants and *Cldn5b* morphants. About 40% of *Cldn5b* morphants displayed hemorrhage at 72 hpf, while *Cldn5a* morphants exhibited no difference with control embryos (Fig.5E and 5F). Interestingly, *Cldn5b* morphants exhibited more pruning structures than control embryos (Fig.5G).

We further observed the status of vasculature in *Cldn5b* morphants. Quantum Dots injection experiment revealed that hemorrhage was obvious in embryos with a lower dosage of *Cldn5b* MO (0.5 pmol), and there was an absence of blood circulation in embryos with a higher dosage of *Cldn5b* MO (1.0 pmol) (Fig.5H). Injection of *Cldn5b* mRNA could partially rescue the hemorrhagic phenotype in *ECAL-1* morphants, while *Cdh5* mRNA could not (Fig.5I). All these results suggested that endothelial tight junction protein Cldn5b functioned downstream of *ECAL-1* to control the cerebrovascular homeostasis, including integrity and pattern formation.

### *ECAL-1* May Protect Cldn5b by Acting As a Sponge of miR-23a

It is known that the function of lncRNAs correlates with their localization within cells (Chen, 2016; Ulitsky and Bartel, 2013). Using fluorescence *in situ* hybridization and PCR quantification of cytoplasmic and nuclear RNA fractions, we identified that *ECAL-1* was expressed in cytoplasm (Fig.6A and 6B), implicating its non-transcriptional modulation on *Cldn5b* expression. Given the canonical action modality of lncRNAs as molecular sponge (Chen, 2016; He et al., 2015), we further conducted bioinformatic prediction. We found that both *ECAL-1* and *Cldn5b* contained a binding site for miR-23a (Fig.6C), which was also widely expressed in the brain (Fig.6E).

**Figure 6.**
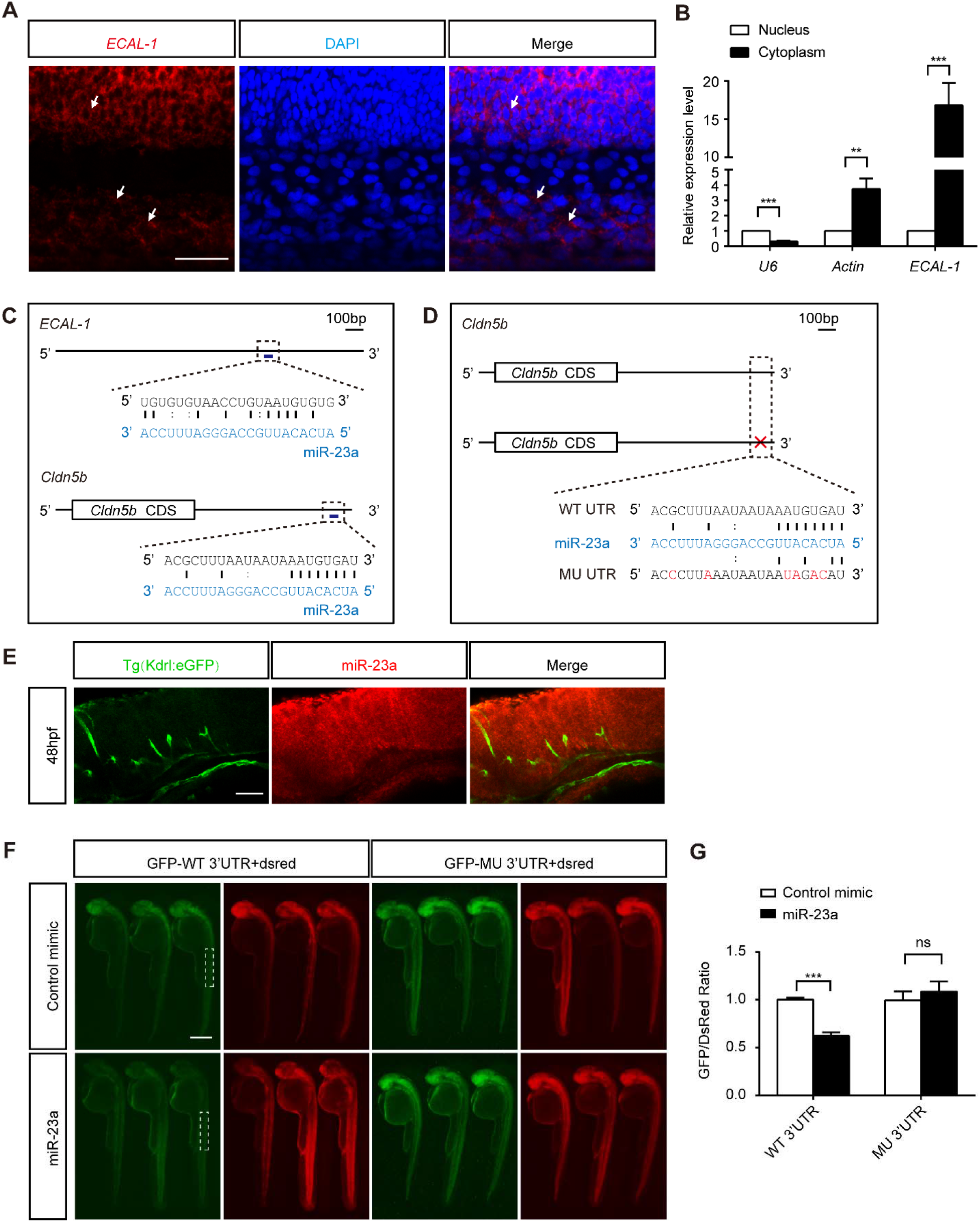
*ECAL-1* promotes Cldn5b by acting as a sponge of miR-23a. **(A)** Fluorescence *in situ* hybridization (FISH) for *ECAL-1* and co-localization with DAPI. **(B)** Quantification of *ECAL-1*, *Actin* and *U6* expression level in nuclear and cytoplasmic fractions of 30hpf embryos. Each group has more than 30 embryos, and this experiment was conducted three times. Statistical analysis was conducted using unpaired Student’s two-tailed *t*-test. **(C)** Diagram of miR-23a target sites in *ECAL-1* and *Cldn5b*. **(D)** Diagram of the 3’UTR of the In vivo reporter assay used. **(E)** Fluorescence *in situ* hybridization (FISH) for miR-23a and immunofluorescence for GFP in hindbrain. **(F)** In vivo reporter assay of the GFP mRNA bearing wild-type *Cldn5b* 3’UTR or mutated 3’UTR co-injected with control mimic, miR-23a mimic at 30 hpf. dsRed mRNA serves as an internal control. **(G)** Quantification of fluorescence density in areas outlined by white dash frame in G. The value of density is calculated by ImageJ. Each group has about 30 embryos, and we only observed the embryos with normal gross morphology, and analyzed representative embryos. This assay was performed three times. Statistical analysis was conducted using unpaired Student’s two-tailed *t*-test. Values are means□±□SEM. **p<0.01; ***p<0.001. Scale bar: 10μm (A); 50μm (E); 500μm (F).

**Figure 7.**
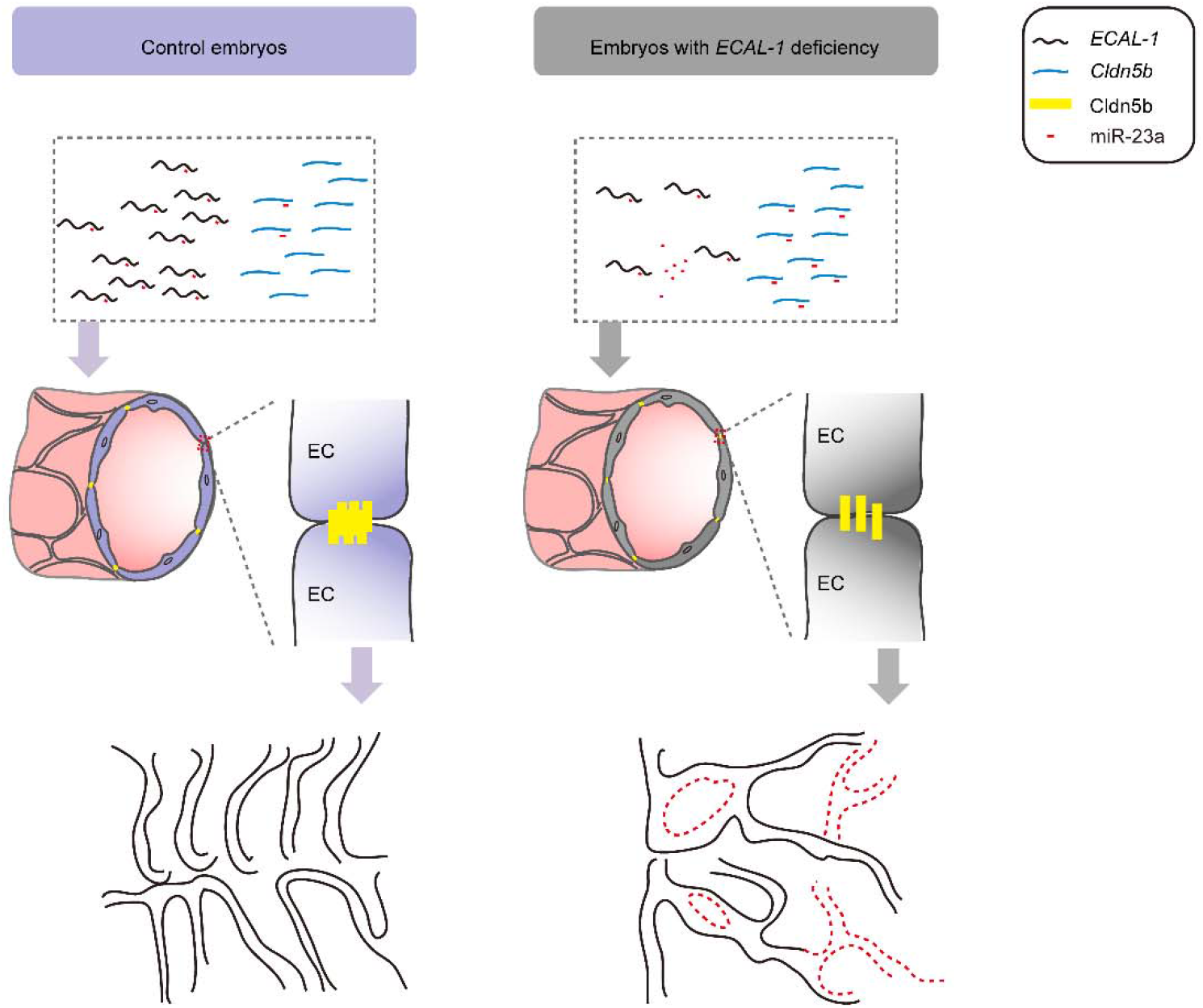
Functional model of *ECAL-1* in cerebrovascular pattern formation. Our work revealed that *ECAL-1* regulated Cldn5b by acting as a sponge of miR-23a during maintaining cerebrovascular pattern. When knockdown of *ECAL-1*, excessive miR-23a inhibit the translation of tight junction protein Cldn5b, resulting in anomalous connection between ECs and abnormal cerebrovascular pattern, even angiorrhexis and blood cells leaked out of vasculature.

To determine whether *Cldn5b* was a target of miR-23a *in vivo*, we co-injected *GFP* mRNAs, which contained wild-type or mutated *Cldn5b* mRNA 3′UTR following GFP coding sequence (CDS), with the mixture containing *dsRed* mRNAs and control or miR-23a mimics at the 1-cell stage. This approach induced a significant downregulation of the expression of GFP bearing a WT 3’UTR, while the expression of GFP bearing a MU 3’UTR was not affected (Fig.6F and 6G), indicating that miR-23a inhibited the translation of *Cldn5b* by targeting Cldn5b 3’UTR. Taken together, *ECAL-1* may promoted Cldn5b expression by acting as a sponge of miR-23a.

## Discussion

In this study, we demonstrated that *ECAL-1* was an essential LncRNA component for cerebrovascular integrity in zebrafish. *ECAL-1* determined cerebrovascular pattern formation through modulating CtAs morphology, EC proliferation and connection. Furthermore, we identified the tight junction protein, *Cldn5b*, as a critical target of *ECAL-1*, which may tether miR-23a to achieve the modulation in cytoplasm.

Increasing evidences suggest that lncRNAs are of great guiding significance for basic biology (Derrien et al., 2012; Klattenhoff et al., 2013). Our results confirmed the spatial-temporal expression of *ECAL-1* in ECs and neurons. To the best of our knowledge, our work for the first time reported the effects of lncRNA on the maintenance of CtAs morphology and cerebrovascular pattern formation. Loss-of-function experiments revealed the indispensability of *ECAL-1* in brain angiogenesis, and its deficiency caused irregularity of cerebrovascular pattern and EC connection. Of note, despite the difference of intracranial hemorrhage rate in morphants, MP mutants and ECAL-1 mutants, it was consistent that defect of *ECAL-1*, rather than its encoded micropeptide, caused aberrant CtAs and EC connection. Potential reasons may underlie the hemorrhagic discrepancy, and the most likely possibility is that genetic compensation resulted in differential phenotype between morphants and Cas9 mutants. Rescue analysis by *ECAL-1* mRNA and micropeptide coding sequence in *ECAL-1* morphants confirmed that hemorrhage were owing to *ECAL-1* absence, and not relevant to any off-target effects or its encoded micropeptide. The evidence that MP mutants without translation of micropeptide exhibited normal cerebral vasculature and no hemorrhage also supported the conclusion.

It is clear that mural cells and junction proteins are pivotal elements for vascular integrity (Dejana et al., 2009; Obermeier et al., 2013; Spadoni et al., 2017). In the present study, no defects of pericyte coverage in *ECAL-1* morphants were observed, whereas ultra-structure analysis revealed the aberrant connection between endothelial cells in *ECAL-1* morphants. These findings precluded the contribution of pericyte coverage in *ECAL-1*-defect-induced cerebrovascular disorders.

The core roles of tight junction protein Claudin5 (CLDN5) in blood-brain barrier permeability of mammals have been identified (Nitta et al., 2003) (He et al., 2015). In zebrafish, Cldn5a expanded the expression in brain vessels from 72 hpf (van Leeuwen et al., 2018), while *Cldn5b* is enriched in brain vasculature at 48 hpf (Xie et al., 2010). *Cldn5a* is involved in brain ventricular development (Zhang et al., 2010), and the function of *Cldn5b* remains unknown. Our work with loss-of-function and rescue experiments revealed that *Cldn5b* dominated the development of cerebrovascular pattern, and controlled vascular integrity. Interestingly, we noticed that the degree of bleeding was more severe in *Cldn5b* morphants than in *ECAL-1* morphants. This inconsistency probably results from that *ECAL-1* is not the only upstream regulator.

The functions of lncRNAs are not only related to its subcellular localization (Chen, 2016; Ulitsky and Bartel, 2013), but also involved in the flank genes. Our data indicated that *ECAL-1* may act as a sponge of miR-23a to regulate the translation of *Cldn5b* in the cytoplasm, and it did not regulate flanking genes expression *in cis* (Fig. S5). Our results further demonstrated that miR-23a inhibited the translation of *Cldn5b*, which was different from the previous report that miR-23a inhibited the EC permeability when its overexpression was conducted in HUVEC and mouse model (Li et al., 2016). This discrepancy maybe attributed to the differential response in different species.

It also deserves to note that neurovascular communication also regulates cerebrovascular development, including vascular integrity (Xu et al., 2017). Herein we identified the neuron-enrichment of *ECAL-1* and miR-23a. Thus, we can’t preclude the possibility that neuronal *ECAL-1-*miR-23a axis indirectly affected cerebrovascular homeostasis.

In summary, our findings reported here provide theoretical basis for underlying mechanism of cerebrovascular homeostasis, and prompt the investigation of lncRNAs in its related diseases.

## Materials and Methods

### Zebrafish Care and Lines

The Tubingen (TU), *Tg(Kdrl:eGFP)*, *Tg(Fli1:neGFP)*^*y7*^ and *Tg(Kdrl:HsHRAS-mCherry)*^*s896*^ zebrafish lines were raised, mated and staged as described previously (Kimmel et al., 1995). Fish maintenance was in accordance with guidelines of the Institutional Review Board of the Institute of Health Sciences, Shanghai Institutes of Biological Sciences, Chinese Academy of Sciences (Shanghai, China). All animal experiments have local approval and all animal experiments were carried out in accordance with the National Institutes of Health Guide for the Use of Laboratory Animals and were approved by the Biological Research Ethics Committee of Institute of Health Sciences.

### Euthanasia of zebrafish

All animal experiments were performed on zebrafish embryos less than 120 hpf and euthanasia was performed by rapid freezing followed by maceration.

### Morpholinos and mRNA Injection

All morpholinos (MOs) were purchased from GeneTools (Philomath, OR) and dissolved in RNA free water (without treatment of DEPC). MOs against *ECAL-1* were designed to block the micropeptide translation initiation or modify the splicing site. A scramble MO was used as a control.

Capped mRNA was synthesized with Sp6 (Promega, WI) or T7 RNA polymerase (Roche, Mannheim). Embryos at one-cell stage were injected with 1 nL MO or/and mRNA, and the concentrations were as follows unless specified elsewhere: *ECAL-1* MO, 1 mmol/L; MP MO, 1 mmol/L; Control MO, 1 mmol/L; *ECAL-1* mRNA, 200 ng/μL; MP mRNA 200 ng/μL, *dsRed* mRNA, 100 ng/μL.

### *In Vitro* Transcription of Cas9 and gRNA

The Cas9 mRNA were synthesized by *in vitro* transcription using T7 RNA polymerase (Roche, Mannheim). The target sites of gRNA starting with “GG” or “GA” were designed manually, and the sequence of a T7 or SP6 promoter was added to the 5’-upstream of gRNA sequence. The Cas9 mRNA and gRNAs were co-injected into 1-cell stage embryos. Each embryo was injected with 1 nL solution containing 200 ng/μL Cas9 mRNA and 100 ng/μL gRNA. The sequence of gRNAs (including T7 promoter and target site) were listed in Table.

### Genotype Identification of Mutant Fish Line

Based on the gRNA target site, we design primers to amplify the wild-type or mutated region. We screened the founders by PCR and sequencing. To further confirmation the genotype of ECAL-1 mutant, we further designed a pair of primers that could only obtain bands in the wild type embryos, not in ECAL-1 mutants. Primers for genotype identification are shown in supplementary.

### O-dianisidine Staining and Scoring of Hemorrhage

TU embryos were collected at 72 hpf, and were stained as previously described (Paffett-Lugassy and Zon, 2005). Embryos with normal gross morphology were enrolled to account the hemorrhage rate, which was divided into four categories, including no hemorrhage, a small hemorrhage site, a large hemorrhage site, and two or more hemorrhage sites.

### RNA Extraction and Quantitative Reverse Transcriptase-Polymerase Chain Reaction (RT-PCR)

The total RNA from embryos was extracted using TRIzol reagent (Invitrogen) according to the protocol. The total RNA extracted was used to generate cDNA by using Super Script II reverse transcriptase with random primer for mRNA. RT-PCR was performed using SYBR Green (TOYOBO, Japan). The relative RNA amount was calculated with the ΔΔCt method and normalized with internal control β-actin.

### Fluorescent-Activated Cell Sorting (FACS)

To isolate GFP-positive endothelial cells from embryos, about 200 *Tg(Kdrl:eGFP)* embryos at 30 hpf were digested into single cell with 1 mL 0.25% trypsin (Gibico, MD) at 28.5 □ until no clumps of tissue are visible to the naked eye. The cells were precipitated by centrifugation (400g, 5min) after adding FBS to stop digestion. Then the cell were rinsed with 10% FBS/PBS for three times. After filtration, the cells resuspended solution (10% FBS/PBS) was sorting by (BD SORP FACSAria).

### Isolation of Cytoplasmic and Nuclear RNA Fractions

About 200 *Tg(Kdrl:eGFP)* embryos at 48hpf were digested into single cell in accordance with 2.8. Then isolation of cytoplasmic and nuclear RNA fractions was performed as previously described (Chen et al., 2008). Briefly, cells pellets were resuspended in lysis buffer (10mM Tris (pH8.0), 140mM NaCl, 1.5mM MgCl2, 0.5% Igepal, 2mM vanadyl robonucleoside complex (VRC, Invitrogen, CA), incubated on ice for 5 min. The lysate was centrifuged at 1000g for 3 min at 4 □ to pellet the nuclei and the supernatant was the cytoplasmic fraction.

### Whole-mount *in situ* Hybridization

Whole-mount *in situ* hybridization with Digoxigenin (Roche, Mannheim) labeled probes was performed in wild type embryos as previously described (Thisse and Thisse, 2008). The sequence of probes for *Kdrl* and *Cmyb* were described previously (Krueger et al., 2011; North et al., 2007). The LNA probe for miR-23a was purchased from Exiqon (Vedbaek, Denmark). The LNA sequence and primers for *in situ* hybridization probes are shown in supplementary.

### Fluorescence *in situ* Hybridization and Immunofluorescence

Fluorescence *in situ* hybridization with Digoxigenin (Roche, Mannheim) labeled probe was performed in wild type embryos at the first and second day as previously described (Thisse and Thisse, 2008). The embryos were blocked with 2% Blocking Reagent (Roche, Mannheim) for 1h, and then incubated in a solution of anti-Digoxigenin-POD Fab Fragments (Roche, Mannheim, 1:1000). The samples were rocked gently overnight at 4 □, and washed 6 times for 20 min each with PBST (0.1% Tween in PBS). A buffer containing Cy3 (PerkinElmer, Waltham, 1:50) was used to stain the samples for 40 min.

For immunofluorescent imaging, the embryos were incubated with a blocking buffer (PBS with 0.1% tween20, 0.1% TritonX100, 10% goat serum, 1% BSA) for 1h, and then with primary antibody overnight at 4 □. After washing six times with PBST, embryos were incubated with a secondary antibody (goat anti-mouse IgG, Alexa 488/594; Invitrogen, CA, 1:1000), and imaging ensued.

Antibodies used in this study: anti-GFP (Yeasen, China, 1:1000); anti-Claudin5 (Invitrogen, CA, 1:100).

### Confocal Imaging and Analysis

Confocal imaging was performed as previously described (Chen et al., 2016; Zou et al., 2011). The embryos were anesthetized in 0.04% Tricaine (Sigma-Aldrich, MO) medium, then were mounted in low melting point Agarose (Sigma-Aldrich, MO). We scanned the interested area with 1.5 μm step size and the format was 1,024×1,024 pixel at 400 Hz. All the confocal images were lateral views, dorsal was up, and anterior to the left unless specifically noted. Both CtAs sprouting in hindbrain and abnormal CtAs were scored in the confocal images.

### *In Vivo* Fluorescence Protein Assay

Modified GFP mRNAs bearing *Cldn5b* 3’UTR (wild-type or mutated) following the GFP open reading frame were synthesized by *in vitro* transcription, and they were separately co-injected with miR-23a/control mimic and *dsRed* mRNA into 1-cell stage embryos. The concentrations of the mRNA and mimics were as follows: GFP mRNA, 50 ng/μL; *dsRed* mRNA, 50 ng/μL; miR-23a/control mimic, 2.5 μM. Fluorescence intensity of GFP and dsRed in trunk were quantified by ImageJ at 30 hpf and dsRed was used as an internal control.

### Angiography

Angiography was performed as previously described (Schmitt et al., 2012). Briefly, 55 hpf *Tg(Kdrl:eGFP)* larvae were anesthetized in 0.04% Tricaine (Sigma-Aldrich, MO) medium, and then 2 nl Quantum Dots were injected into venous sinus through a microinjection setup with glass capillaries.

### Bioinformatic Analysis

The online tools of genie.weizmann.ac.il/pubs/mir07/mir07_prediction.html and TargetScan were used to identify potential binding sites of miR-23a in *ECAL-1* and *Cldn5b* 3’UTR, respectively.

### Statistical Analysis

All the results were generated from at least three independent experiments, and data were presented as means ± SEM. Statistical analysis was performed using GraphPad. Analysis of differences between two groups was conducted the unpaired Student’s two-tailed *t*-test. When more than two groups, statistical differences were performed one-way ANOVA with Tukey’s *post-hoc* test. Differences were considered significant when P < 0.05. Probability values are indicated by * (P < 0.05), ** (P < 0.01), or *** (P < 0.001).

## Acknowledgements

We thank Dr. Didier Stainier (Max Planck Institute, Bad Nauheim, Germany) for providing *Tg(Kdrl:HsHRAS-mCherry)*^*s896*^ transgenic lines; and Dr. Nathan Lawson (University of Massachusetts Medical School, Worcester, USA) for kindly providing the *Tg(Kdrl:eGFP)*. We thank Dr. Jun Li (Shanghai General Hospital, China) for modifying the manuscript; Dr. Jiu-Lin Du (Institute of neuroscience, CAS, Shanghai, China) for kindly providing *Tg(Fli1:neGFP)*; and Dr Jing-Wei Xiong (Peking University, Beijing, China) for providing the plasmids of CRISPR/Cas9 system. We thank all members in Dr. Jing’s lab for helpful discussions and comments on this article. We thank Min Deng for technical assistance and Sheng-Rong Yang for zebrafish husbandry.

## Data availability

All relevant data are within the paper and its Supporting Information files

